# Disruption of Herpes Simplex Virus Type 2 pUL21 Phosphorylation Impairs Secondary Envelopment of Cytoplasmic Nucleocapsids

**DOI:** 10.1101/2024.04.10.588913

**Authors:** Renée L. Finnen, Jamil H. Muradov, Valerie Le Sage, Bruce W. Banfield

## Abstract

The multifunctional tegument protein pUL21 of HSV-2 is phosphorylated in infected cells. We have identified two residues in the unstructured linker region of pUL21, serine 251 and serine 253, as sites of phosphorylation. Both phosphorylation sites are absent in HSV-1 pUL21, which likely explains why phosphorylated pUL21 was not detected in cells infected with HSV-1. Cells infected with HSV-2 strain 186 viruses deficient in pUL21 phosphorylation exhibited reductions in both cell-cell spread of virus infection and virus replication. Defects in secondary envelopment of cytoplasmic nucleocapsids were also observed in cells infected with viruses deficient in pUL21 phosphorylation as well as in cells infected with multiple strains of HSV-2 and HSV-1 deleted for pUL21. These results confirm a role for HSV pUL21 in the secondary envelopment of cytoplasmic nucleocapsids and indicate that phosphorylation of HSV-2 pUL21 is required for this activity. Phosphorylation of pUL21 was not detected in cells infected with HSV-2 strain 186 mutants lacking the viral serine/threonine kinase pUL13, indicating a requirement for pUL13 in pUL21 phosphorylation.

**Importance:** It is well known that post-translational modification of proteins by phosphorylation can regulate protein function. Here, we determined that phosphorylation of the multifunctional HSV-2 tegument protein pUL21 requires the viral serine/threonine kinase pUL13. Additionally, we identified serine residues within HSV-2 pUL21 that can be phosphorylated. Phenotypic analysis of mutant HSV-2 strains with deficiencies in pUL21 phosphorylation revealed reductions in both cell-cell spread of virus infection and virus replication. Deficiencies in pUL21 phosphorylation also compromised secondary envelopment of cytoplasmic nucleocapsids, a critical final step in the maturation of all herpes virions. Unlike HSV-2 pUL21, phosphorylation of HSV-1 pUL21 was not detected. This fundamental difference between HSV-2 and HSV-1 may underlie our previous observations that the requirements for pUL21 differ between HSV species.

## Introduction

The capsid-associated tegument component pUL21 is a multifunctional protein unique to the alphaherpesvirus subfamily of the *Herpesviridae*. Known functions of pUL21 include facilitating transport of capsids along microtubules (1–3), promoting secondary envelopment of capsids (4), regulating nuclear egress of capsids (5, 6), directing protein phosphatase 1 alpha (PP1α) to specific substrates for dephosphorylation (7, 8), triggering autophagic degradation of cyclic GMP-AMP synthase (cGAS) (9), enabling retention of viral genomes within capsids (10) and, most recently, preventing capsids from docking to nuclear pores after nuclear egress (11). With so many important roles played by pUL21 during viral infection, it is not surprising that viruses deficient in pUL21 display a small plaque phenotype and reduced viral replication in comparison to wild type (WT) virus (1, 4, 8, 12–15). The defects in cell-cell spread of virus infection and replication of pUL21 deficient viruses are more pronounced in herpes simplex virus type 2 (HSV-2) than they are in herpes simplex virus type 1 (HSV-1) even though the pUL21s of these species are about 84% identical (13).

In our characterization of the role played by HSV-2 pUL21 in nuclear egress, we utilized phos-tag electrophoresis to demonstrate that pUL21 can modulate phosphorylation of the nuclear egress complex (NEC) proteins, pUL31 and pUL34, by the viral serine/threonine kinase pUs3 (6). During these studies, we detected two forms of HSV-2 pUL21 when infected cell lysates were examined by phos-tag electrophoresis and western blotting with pUL21-specific polyclonal antiserum. The more slowly migrating form of pUL21 was not detected when the infected cell lysate was treated with lambda protein phosphatase (λPP), indicating that pUL21 was phosphorylated. As phosphorylation of a protein often affects its function (16, 17), we wished to further investigate pUL21 phosphorylation.

The goals of this work were first, to gain an understanding of the phosphorylation profile of HSV-2 pUL21 and second, to use this understanding to construct viruses that would allow us to begin assessing phenotypes associated with disruption of pUL21 phosphorylation. We present evidence that disruption of pUL21 phosphorylation impairs cell-cell spread of virus infection as well as viral replication. While disruption of pUL21 phosphorylation did not have an impact on nuclear egress of HSV-2 capsids, secondary envelopment of cytoplasmic HSV-2 nucleocapsids was compromised. We also present evidence that pUL21 phosphorylation requires the viral serine/threonine kinase pUL13.

## Materials and Methods

### Cells and viruses

African green monkey kidney cells (Vero), HeLa cells, human keratinocytes (HaCaT), 293T cells, rabbit kidney cells (RK13) and mouse L cells were maintained in Dulbecco’s modified Eagle medium (DMEM) supplemented with 10% fetal bovine serum (FBS) in a 5% CO_2_ environment. RK13 cells that stably produce HSV-2 pUL21-EGFP (referred to herein as 21-GFP cells) were isolated by transfecting cells with a plasmid encoding UL21-EGFP (1), selecting for G418 resistance, and screening for green fluorescence. L cells that stably produce HSV-2 pUL21 (L21) were isolated by retroviral transduction and selected for puromycin resistance as described previously (1). HSV-2 strains 186 and SD90e were acquired from Dr. Y. Kawaguchi (University of Tokyo) and Dr. D. M. Knipe (Harvard University), respectively; HSV-1 strains F and KOS were acquired from Dr. L. W. Enquist (Princeton University). HSV-2 strain 186 virus lacking pUL21 (ΔUL21) was previously described (1). HSV-2 strain 186 viruses lacking pUL13 were constructed by CRISPR/Cas9 mutagenesis using UL13-specific guide RNAs (Table 1), using methodologies described previously (13). HSV-2 strain 186 viruses carrying mutations in pUL21 were constructed by two-step Red-mediated mutagenesis (18) in *Escherichia coli* strain GS1783, a gift from Dr. G. A. Smith (Northwestern University) using pYEbac373, a recombinant bacterial artificial chromosome (BAC) of the WT 186 genome (1), or pYEbac373_S253A, in which serine 253 of pUL21 was changed to alanine. Primers (Table 2) were used to amplify a PCR product from pEP-Kan-S2, a gift from Dr. N. Osterrieder (Freie Universität Berlin), to change serine 253 of pUL21 to either alanine (S253A) or glutamic acid (S253E), to change serine 251 of pUL21 to alanine (S251A), and to change both serines 251 and 253 to alanine (S251/253A). Primers (Table 2) were used to amplify a PCR product from pEP-mCherry-in, a gift from Dr. G. A. Smith (Northwestern University), to fuse mCherry to the carboxy-terminus of WT, S253A or S253E pUL21. Restriction fragment length polymorphism analysis was used to confirm the integrity of each BAC clone in comparison to WT BAC by digestion with *Eco*RI. Additionally, PCR was used to amplify products spanning UL21 or UL21-mCherry and these products were sequenced to confirm the presence of the intended point mutations, to ensure that no unintended mutations were introduced into UL21 or the mCherry gene and to confirm that the mCherry gene was in frame with UL21. Infectious virus was reconstituted from BAC DNA as described previously (1). Times post infection, reported as hours post infection (hpi), refer to the time elapsed following medium replacement after an inoculation period of one hour.

**Table 1.**
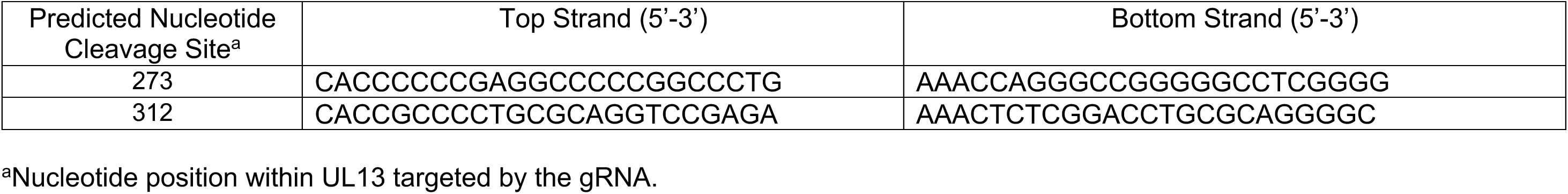
Oligonucleotides used to produce HSV-2 UL13 gRNAs.

**Table 2.**
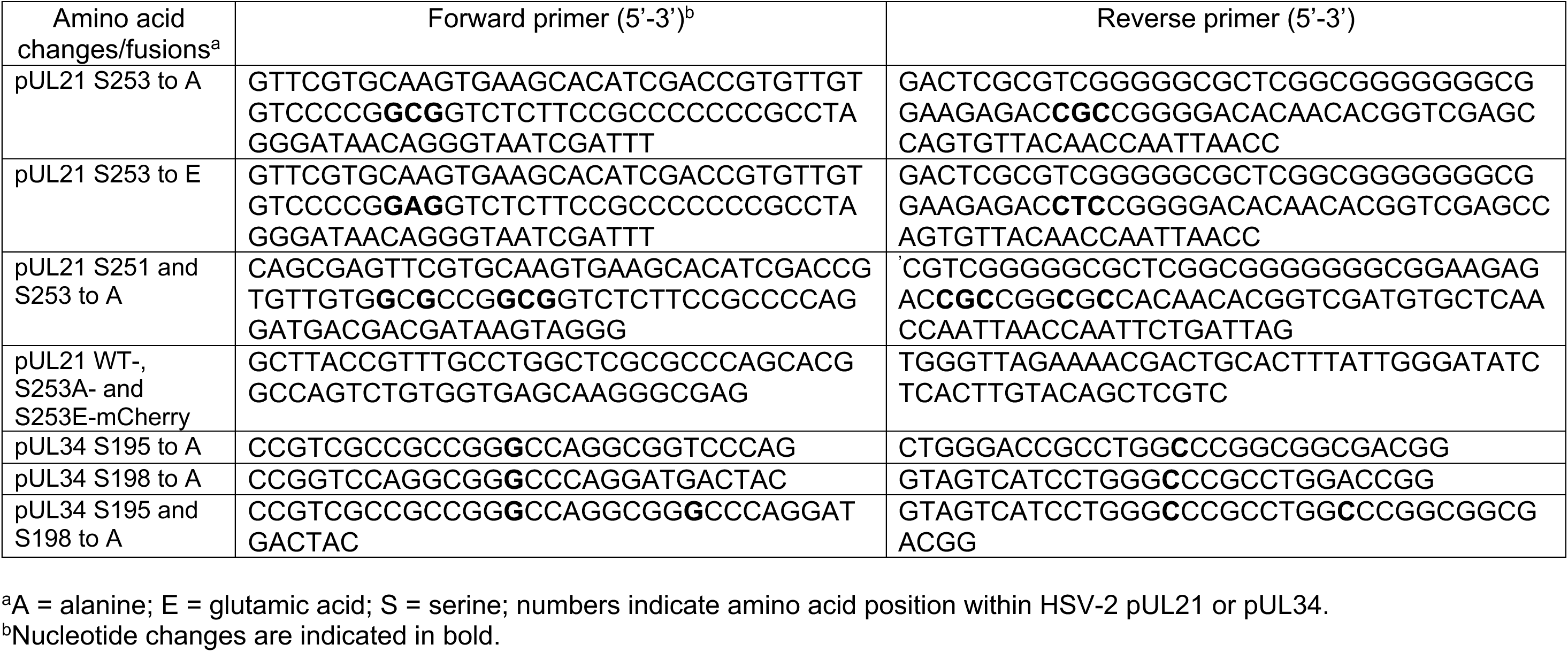
Oligonucleotide primers used for mutagenesis of pUL21 and pUL34.

### Virus replication analysis

For replication analysis, 6 well dishes containing 3 × 10^6^ Vero cells per well were infected at a multiplicity of infection (MOI) of 0.1. All infections were performed in triplicate. After an inoculation period of one hour, cells were treated for 3 min at 37^◦^C with low pH citrate buffer (40 mM Na citrate, 10 mM KCl, 0.8% NaCl) to inactivate extracellular virions, washed once with medium, then incubated at 37^◦^C. Medium and infected cells were harvested together at 48 hpi. Samples were subjected to two freeze/thaw cycles, sonicated briefly in a chilled cup-horn sonicator, and cellular debris was removed from the lysate by brief centrifugation at 1,620 x g in a microcentrifuge. Virus titres were determined in duplicate by plaque assay on L21 cells.

### Plasmid construction and transfection

The construction of plasmids encoding HSV-2 UL21 and UL21-GFP were described previously (1). The construction of plasmids encoding HSV-2 WT Us3 and a Us3 mutant in which the codon for aspartic acid 305 was changed to alanine to disrupt the kinase activity of pUs3, were described previously (19). The construction of a plasmid encoding HSV-2 UL34 was described previously (6). To change serines 195 and 198 of pUL34 to alanine, alone, or in combination, the primers listed in Table 2 were used in successive mutagenic PCR reactions. The PCR-amplified portion of all plasmids was sequenced to ensure that no unintended mutations were introduced. For preparing cellular extracts from transfected cells for western blot analyses, plasmids were transfected into HeLa cells using X-tremeGENE HP DNA transfection reagent (Roche, Laval, QC) according to manufacturer’s protocols.

### Immunological reagents

Rat polyclonal antiserum against HSV-2 pUL21 (1) was used for western blotting at a dilution of 1:600; chicken polyclonal antiserum against HSV-2 pUL34 (5) was used for western blotting at a dilution of 1:500; rat polyclonal antiserum against HSV-2 pUL31 (5) was used for western blotting at a dilution of 1:200; mouse monoclonal antibody against HSV pUL13 (20), a gift from Dr. Y. Kawaguchi (University of Tokyo), was used for western blotting at a dilution of 1:1,000; and mouse monoclonal antibody against β– actin (Sigma, St. Louis, MO) was used for western blotting at a dilution of 1:2,000. Horseradish peroxidase-conjugated rabbit anti-rat IgG and horseradish peroxidase-conjugated goat anti-chicken IgY (Sigma, St. Louis, MO) were used for western blotting at a dilution of 1:80,000. Horseradish peroxidase-conjugated goat anti-mouse IgG (Jackson ImmunoResearch, West Grove, PA) was used for western blotting at a dilution of 1:30,000. Rat polyclonal antiserum against HSV-2 pUs3 (19) was used for indirect immunofluorescence microscopy at a dilution of 1:500.

### Preparation and analysis of cellular protein extracts

To prepare cellular extracts of infected cells for standard western blot analyses, 3 × 10^6^ cells infected at a MOI of 3 or mock infected were washed at 18 hpi with cold phosphate buffered saline (PBS), scraped into 100 μl of PBS containing protease inhibitors (Roche, Laval, QC), then transferred to a microcentrifuge tube containing 50 μl of 3X SDS-PAGE loading buffer. The lysates were repeatedly passed through 28 ½-gauge needles to reduce their viscosity. To prepare cellular extracts of infected or transfected cells for phos-tag western blot analyses, 3 × 10^6^ cells infected at a MOI of 3 or 3 × 10^6^ transfected cells were washed 18 hours after infection or transfection with cold PBS and then scraped into 200 μl cold RIPA buffer (50 mM Tris-HCl pH 7.5, 150 mM NaCl, 0.5% sodium deoxycholate, 1% NP-40, 1% SDS) containing protease inhibitors (Roche, Laval, QC). The lysates were repeatedly passed through 28 ½-gauge needles to reduce their viscosity and 250 units of benzonase nuclease (Santa Cruz Biotechnology, Dallas, TX) were added to each lysate. For lambda protein phosphatase (λPP) treatment, 39 μl of each benzonase-treated lysate was combined with 5 μl of 10X NEB buffer for Protein MetalloPhosphatases (New England Biolabs, Ipswich, MA), 5 μl of 10 mM MnCl_2_ and 1 μl (400 units) of λPP (New England Biolabs, Ipswich, MA). Untreated lysates received 1 μl of H_2_O in place of λPP. The dephosphorylation reactions were carried out for 30 minutes at 30°C. The dephosphorylated lysates were centrifuged for 10 minutes at 14,000 x g and 44 μl of supernatants were transferred to 1.5 mL tubes containing 22 μl 3X SDS-PAGE loading buffer. For western blot analysis, 10 to 20 μl of boiled extract was electrophoresed through regular SDS-PA gels or SDS-PA gels containing 50 μM phos-tag reagent (Fujifilm, Richmond, USA) supplemented with 0.1 mM MnCl_2_. Separated proteins were transferred to PVDF membranes (Millipore, Billerica, MA) and probed with appropriate dilutions of primary antibody followed by appropriate dilutions of horseradish peroxidase-conjugated secondary antibody. The membranes were treated with Pierce ECL western blotting substrate (Thermo Scientific, Rockford, IL) and exposed to film.

### Immunoprecipitation of pUL21-EGFP and pUL21-mCherry and mass spectrometry analysis

Lysates from 21-GFP cells grown on two 150 mm dishes and infected with HSV-2 strain 186 at an MOI of 3 or uninfected were prepared at 18 hpi and immobilized on GFP-TRAP beads (Chromotek, Rosemont, IL) according to manufacturer’s instructions. Lysates from HaCaT cells grown on two 150 mm dishes and infected with HSV-2 strain 186 viruses carrying pUL21-mCherry fusion proteins at an MOI of 3 were prepared at 18 hpi and immobilized on RFP-TRAP beads (Chromotek, Rosemont, IL) according to manufacturer’s instructions. Washed beads were boiled in 2X SDS-PAGE loading buffer to elute the bound proteins, immunoprecipitated proteins were electrophoresed 5 mm into a regular 10% SDS-PA gel, stained with SimplyBlue SafeStain (Invitrogen, Carlsbad, CA) according to manufacturer’s instructions, and then 5 mm X 5 mm slices from the gel were excised. Orbitrap/orbitrap tandem mass spectrometry on proteins eluted from gel slices and digested with trypsin was performed by the Southern Alberta Mass Spectrometry Facility at the University of Calgary. Tandem mass spectra data were searched against a rabbit protein database, a human protein database containing WT HSV proteins, and an HSV database that was also customized for substitution mutations in HSV-2 S253 using Mascot version 2.7.0 (Matrix Science, London, UK), assuming the digestion enzyme trypsin. Mascot was searched with a fragment ion mass tolerance of 0.020 Da and a parent ion tolerance of 10.0 PPM. Oxidation of methionine and phosphorylation of serine, threonine and tyrosine were specified in Mascot as variable modifications. Scaffold version 5.1.0 (Proteome Software Inc., Portland OR) was used to validate peptide and protein identifications. Peptide identifications were accepted if they could be established at greater than 95.0% probability by the Peptide Prophet algorithm (21) with Scaffold delta-mass correction. Protein identifications were accepted if they could be established at greater than 95.0% probability and contained at least one identified peptide. Protein probabilities were assigned by the Protein Prophet algorithm (22). Proteins that contained similar peptides and could not be differentiated based on MS/MS analysis alone were grouped to satisfy the principles of parsimony. The identity of the immunoprecipitated proteins that were predicted to be phosphorylated in our mass spectrometry analyses can be found in the supplementary data.

### Transmission electron microscopy (TEM)

Vero cells growing in 100 mm dishes were infected at a multiplicity of infection of 3 and processed for TEM at 18 hpi. Infected cells were washed 3 times with PBS before fixing in 1.5 ml of 2.5% electron microscopy-grade glutaraldehyde (Ted Pella, Redding, CA) in 0.1 M sodium cacodylate buffer (pH 7.4) for 60 minutes. Cells were collected by scraping into fixative and pelleted by repeated centrifugation at 300 x g for 5 minutes. Cell pellets were carefully enrobed in an equal volume of molten 5% low-melting temperature agarose (Lonza, Rockland, ME) and allowed to cool. Specimens in agarose were incubated in 2.5% glutaraldehyde in 0.1 M sodium cacodylate buffer (pH 7.4) for 90 minutes and postfixed in 1% osmium tetroxide for 60 minutes. The fixed cells in agarose were rinsed 3 times with distilled water and stained in 0.5% uranyl acetate overnight before dehydration in ascending grades of ethanol (30 to 100%). Samples were transitioned from ethanol to infiltration with propylene oxide and embedded in Embed-812 hard resin (Electron Microscopy Sciences, Hatfield, PA). Blocks were sectioned at 50 to 60 ριm and stained with uranyl acetate and Reynolds’ lead citrate. Images were acquired using a Hitachi H-7000 transmission electron microscope. At least 9 independent micrographs of cells infected with each virus strain were scored to determine the ratio of non-enveloped to enveloped cytoplasmic nucleocapsids.

## Results

### Phosphorylation of pUL21 is a conserved property of HSV-2 but not of HSV-1

Our recent studies on the regulation of phosphorylation of the HSV-2 NEC by pUL21 revealed that approximately half of the pUL21 in Vero cells infected with HSV-2 strain 186 was phosphorylated (6). To examine whether pUL21 is phosphorylated in cells infected with other HSV-2 and HSV-1 strains, cells were infected with HSV-2 strains 186 and SD90e and HSV-1 strains F and KOS and infected cell extracts were subjected to phos-tag SDS-PAGE and western blot analysis with pUL21 antiserum. As can be seen in Figure 1A, both phosphorylated and non-phosphorylated forms of pUL21 were detected in HSV-2 infections but only non-phosphorylated pUL21 was detected in HSV-1 infections. Thus, phosphorylation of pUL21 appears to be a conserved property of HSV-2 but not of HSV-1. The cells used for the experiment depicted in Figure 1A were HeLa cells and similar results were also observed with infected Vero and HaCaT cells (Figure 1B). Thus, phosphorylation of HSV-2 pUL21 and lack of phosphorylation of HSV-1 pUL21 is not cell-type dependent.

**Figure 1.**
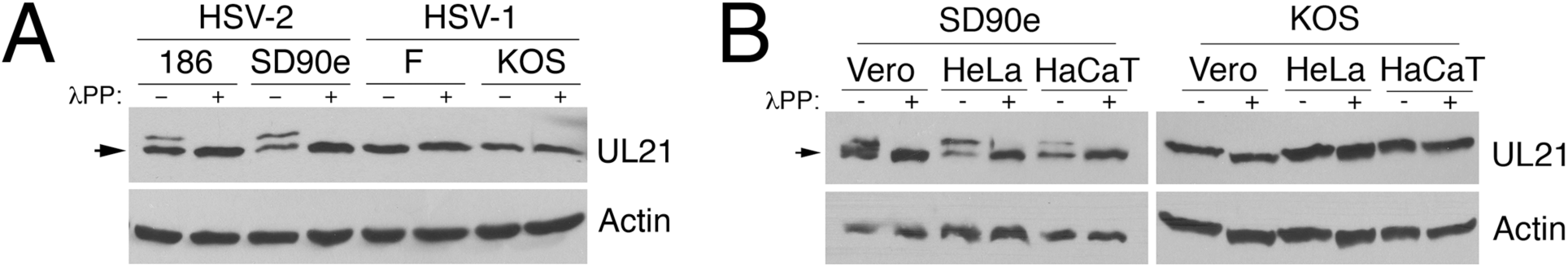
Phosphorylation of pUL21 differs between HSV-2 and HSV-1. Whole cell lysates prepared at 18 hours post infection from HeLa cells (**A**) or the indicated cell types (**B**) infected with the indicated viruses were either treated with lambda protein phosphatase (λPP) or untreated then electrophoresed through phos-tag SDS polyacrylamide gels and proteins were transferred to PVDF membranes. Membranes were probed with antisera indicated on the right of each panel. Arrows denote the position of unphosphorylated pUL21.

### Phosphorylation of pUL21 is not detected in cells infected with HSV-2 strains deficient in pUL13

Phosphorylation of pUL21 was not detected in cells transfected with an HSV-2 pUL21 expression construct (Figure 2A), suggesting that pUL21 is only phosphorylated in infected cells. These observations narrow down the kinase responsible for phosphorylating HSV-2 pUL21 to a virally encoded kinase or a cellular kinase activated upon infection. The responsible kinase is likely not the viral serine-threonine kinase pUs3 as phosphorylated pUL21 was still detected following infection with an HSV-2 Us3 null mutant (6). Another candidate kinase is the viral serine-threonine kinase pUL13 (23–27). To investigate the ability of pUL13 to phosphorylate pUL21, we used CRISPR/Cas9 mutagenesis to knock out UL13 from HSV-2 strain 186 (ΔUL13). A comparison of the UL13 genes from two independent ΔUL13 viruses that were recovered, designated ΔUL13-3 and ΔUL13-6, with WT UL13 is depicted in Figure 2B. The stop codon of the UL14 open reading frame, which resides within the UL13 open reading frame (28, 29), was left intact in both viruses. Both viruses contain a frameshift in UL13 after codon 96 that results in the production of a truncated pUL13 with non-pUL13 amino acids on the carboxy-terminus; 193 and 194 amino acids in the case of ΔUL13-3 and ΔUL13-6, respectively. Both viruses yielded smaller plaques in comparison to WT virus (Figure 2C), as has been previously reported for HSV-2 strain 186 lacking pUL13 (20), and pUL13 was not detected in infected cell lysates (Figure 2D). Phosphorylated pUL21 was not detected in infected cell lysates from both pUL13 deficient viruses (Figure 2E). The phosphorylation pattern of the HSV-2 NEC component pUL34, which can be phosphorylated by pUs3 on serines 195 and 198 (6), was unaffected by the absence of pUL13 (Figure 2E). The designation of the phosphoforms of pUL34 in Figure 2E was determined by phos-tag western analysis of extracts from HeLa cells transfected with a WT pUL34 expression construct or pUL34 expression constructs in which serine 195 and/or 198 were changed to alanine along with WT or kinase-dead (KD) Us3 expression constructs (Figure 2F). As the phosphoform of pUL21 is specifically lacking in the absence of pUL13, this makes pUL13 a strong candidate for the kinase that phosphorylates pUL21.

**Figure 2.**
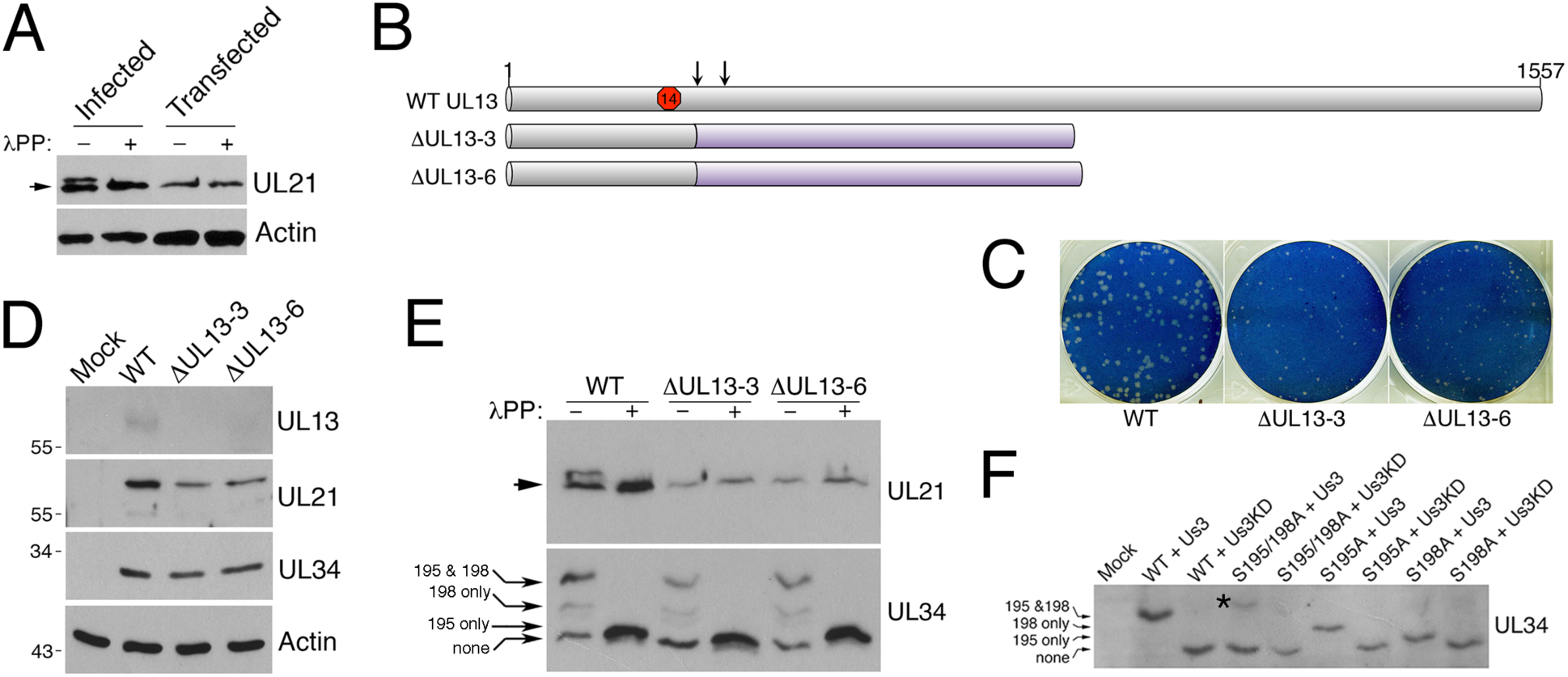
Phosphorylation of pUL21 is not detected in the absence of pUL13. **A.** Whole cell lysates prepared at 18 hours post infection/transfection from HeLa cells infected with HSV-2 strain 186 or transfected with HSV-2 pUL21 expression plasmid were either treated with lambda protein phosphatase (λPP) or untreated then electrophoresed through phos-tag SDS polyacrylamide gels and proteins were transferred to PVDF membranes. Membranes were probed with antisera indicated on the right of each panel. Arrow denotes the position of unphosphorylated pUL21. **B.** Deletions introduced into UL13 by CRISPR/Cas9 mutagenesis. Grey bars indicate wild type (WT) UL13 sequence; purple bars indicate non-pUL13 coding sequence arising from the introduced frameshift mutations; red octagon indicates the stop codon for UL14; vertical arrows indicate gRNA-directed cleavage sites within UL13. **C.** Vero cell monolayers were infected with the indicated viruses, overlayed with medium containing 1% methocel then fixed and stained with 0.5% methylene blue in 70% methanol at 3 days post infection to visualize plaque formation. **D.** Whole cell lysates prepared at 18 hours post infection from HeLa cells infected with the indicated viruses or mock infected were electrophoresed through standard SDS polyacrylamide gels and proteins were transferred to PVDF membranes. Membranes were probed with antisera indicated on the right of each panel. Migration positions of molecular weight markers are indicated on the left of each panel. **E.** Whole cell lysates prepared at 18 hours post infection from HeLa cells infected with the indicated viruses were either treated with lambda protein phosphatase (λPP) or untreated then electrophoresed through phos-tag SDS polyacrylamide gels and proteins were transferred to PVDF membranes. Membranes were probed with antisera indicated on the right of each panel. Arrow denotes the position of unphosphorylated pUL21. Migration positions of phosphoforms of pUL34 are indicated on the left of the bottom panel by the long arrows. **F.** HeLa cells were mock transfected or co-transfected with the pUL34 and pUs3 expression plasmids indicated above each lane. Whole cell extracts prepared at 18 hours post transfection were electrophoresed through a phos-tag SDS polyacrylamide gel, proteins were transferred to a PVDF membrane and the membrane was probed with HSV-2 pUL34 antiserum. Migration positions of phosphoforms of pUL34 are indicated on the left of the panel by the long arrows. Asterisk indicates a phosphoform of pUL34 detected in the absence of serines 195 and 198, which may have arisen from a cryptic phosphorylation site recognized by pUs3 in the absence of bona fide substrates.

### Determination of HSV-2 pUL21 phosphorylation sites

To identify the phosphorylation site(s) in pUL21, pUL21-EGFP was immunoprecipitated from infected or uninfected 21-GFP cells and immunoprecipitated proteins were analyzed by mass spectrometry. This analysis predicted serine 253, located in the unstructured linker region of pUL21 that connects its structured amino- and carboxy-terminal domains (30, 31), as the sole phosphorylation site. As shown in Figure 3A, the corresponding amino acid in HSV-1 pUL21 is glycine, providing a plausible explanation for the lack of pUL21 phosphorylation in HSV-1 infected cells (Figure 1A). In keeping with our evidence in support of pUL13 as the kinase that phosphorylates pUL21 (Figure 2E), phosphorylation of pUL21-EGFP was only detected in proteins immunoprecipitated from infected 21-GFP cells.

**Figure 3.**
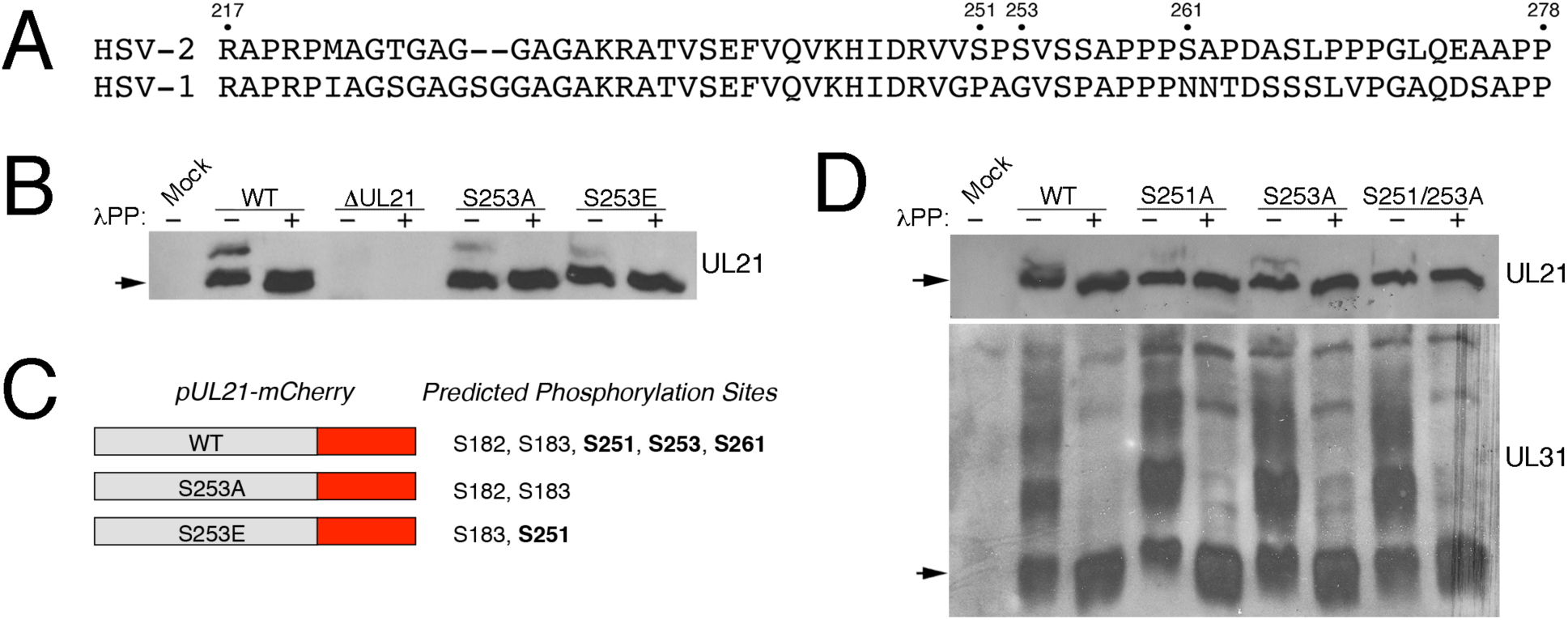
Determination of pUL21 phosphorylation sites. **A.** Alignment of the pUL21 linker regions of HSV-2 and HSV-1. The coordinates of the linker region were defined for HSV-1 pUL21 by Graham and colleagues (8). Numbers above the alignment refer to HSV-2 pUL21. **B, D.** Whole cell lysates prepared at 18 hours post infection from HeLa cells infected with the indicated viruses were either treated with lambda protein phosphatase (λPP) or untreated then electrophoresed through phos-tag SDS polyacrylamide gels and proteins were transferred to PVDF membranes. Membranes were probed with antisera indicated on the right of each panel. Arrows denote the position of unphosphorylated proteins. **C.** Phosphorylation sites predicted in pUL21-mCherry. Sites in bold reside in the linker region; all are not conserved in HSV-1 pUL21 (see alignment in panel **A**). Sites in the amino-terminal domain are conserved in HSV-1 pUL21.

To validate the prediction of serine 253 as the site of HSV-2 pUL21 phosphorylation, we constructed HSV-2 strain 186 viruses carrying alanine (S253A) or glutamic acid (S253E) substitution of serine 253 using *en passant* mutagenesis (18). When we examined the phosphorylation status of pUL21 in cells infected with viruses carrying mutations in serine 253, some phosphorylation of pUL21 was still detected, however, the level of pUL21 phosphorylation appeared to be diminished in comparison to cells infected with WT virus (Figure 3B).

The diminished level of pUL21 phosphorylation in cells infected with either S253A or S253E supports the prediction of serine 253 as a site of pUL21 phosphorylation, however, there may be an additional phosphorylation site that was not detected in our initial mass spectrometry analysis or, alternatively, a cryptic phosphorylation site that is used in the absence of serine 253. To decipher the full spectrum of possible HSV-2 pUL21 phosphorylation sites, we repeated the mass spectrometry analysis of pUL21-derived peptides using immunoprecipitated proteins purified from HaCaT cells infected with viruses carrying WT, S253A and S253E pUL21 with mCherry fused to the carboxy-terminus (WT-mCh, S253A-mCh and S253E-mCh). This analysis predicted two additional phosphorylation sites in the linker region of HSV-2 pUL21, serines 251 and 261, as well as two phosphorylation sites in the structured amino-terminus of HSV-2 pUL21, serines 182 and 183 (Figure 3C). The additional predicted phosphorylation sites in the linker region are also not conserved in HSV-1 pUL21 (Figure 3A) while the predicted phosphorylation sites in the amino-terminus are conserved in HSV-1 pUL21. These observations coupled with the observation that phosphorylation of sites in the linker region were more prevalent than those in the amino-terminus (Table 3) focused our follow-up efforts on the predicted phosphorylation sites in the linker region of HSV-2 pUL21. The data shown in Table 3 indicated that serine 251 was the most prevalent phosphorylation site of HSV-2 pUL21 in this mass spectrometry analysis. We reasoned that changing serine 251 to alanine (S251A) and/or changing both serines 251 and 253 to alanine (S251/253A) might yield viruses with even greater reductions in pUL21 phosphorylation than observed in S253A infections and, if so, that these viruses would be valuable tools for investigating phenotypes associated with disruption of pUL21 phosphorylation. The level of pUL21 phosphorylation was further reduced in cells infected with S251A in comparison to cells infected with S253A while phosphorylated pUL21 was not detected in cells infected with S251/253A (Figure 3D). Thus, both serine 251 and serine 253 are sites of phosphorylation on HSV-2 pUL21.

**Table 3.** Analysis of phosphorylation site prevalence in HSV-2 pUL21.

| Source of pUL21 peptides – database<sup>a</sup> | Phosphorylation on S182/total<sup>b</sup> | Phosphorylation on S183/total | Phosphorylation on S251/total | Phosphorylation on S253/total | Phosphorylation on S261/total |
| --- | --- | --- | --- | --- | --- |
| WT-mCh – human + HSV | 1/39 | 2/39 | 1/12 | 1/12 | 1/12 |
| WT-mCh – HSV | 1/33 | 1/33 | 1/8 | 0/8 | 0/8 |
| S253A-mCh – human + HSV | 1/38 | 2/38 | ND<sup>c</sup> | NA<sup>d</sup> | ND |
| S253A-mCh – HSV | 1/26 | 2/26 | 0/2 | NA | 0/2 |
| S253E-mCh – human + HSV | 0/35 | 2/35 | ND | NA | ND |
| S253E-mCh – HSV | 0/30 | 1/30 | 2/3 | NA | 0/3 |

<sup>a</sup>Raw data were analyzed against a human database containing WT HSV protein sequences or a database containing HSV protein sequences with WT or customized pUL21 sequences; <sup>b</sup>Total refers to the total number of peptides identified that encompassed this particular serine; <sup>c</sup>ND = not done as customized pUL21 sequences were not part of the human + HSV database; <sup>d</sup>NA = non-applicable as S253 was not present in the source peptides.

### Diminished pUL21 phosphorylation impairs both cell-cell spread of virus infection and virus replication

While titrating stocks of HSV-2 viruses with defects in pUL21 phosphorylation, we noticed that plaques formed by these viruses on Vero cells were much smaller than plaques formed by WT HSV-2, suggesting that diminished pUL21 phosphorylation impairs cell-cell spread of virus infection. To quantify this, we performed plaque size analysis on WT, S251A, S253A, S253E, S251/253A and τιUL21 viruses in Vero cells. Plaques formed by all viruses with defects in pUL21 phosphorylation were significantly smaller than plaques formed by WT virus but larger than virus lacking pUL21 (Figures 4A, B). The small plaque phenotype exhibited by all viruses with defects in pUL21 phosphorylation was complemented by providing pUL21 *in trans* (Figure 4C) confirming that this phenotype can be ascribed to the mutations introduced into pUL21. The replication of viruses with defects in pUL21 phosphorylation, assessed by endpoint titration at 48 hpi following low multiplicity infection of Vero cells, was also significantly diminished in comparison to WT (Figure 4D).

**Figure 4.**
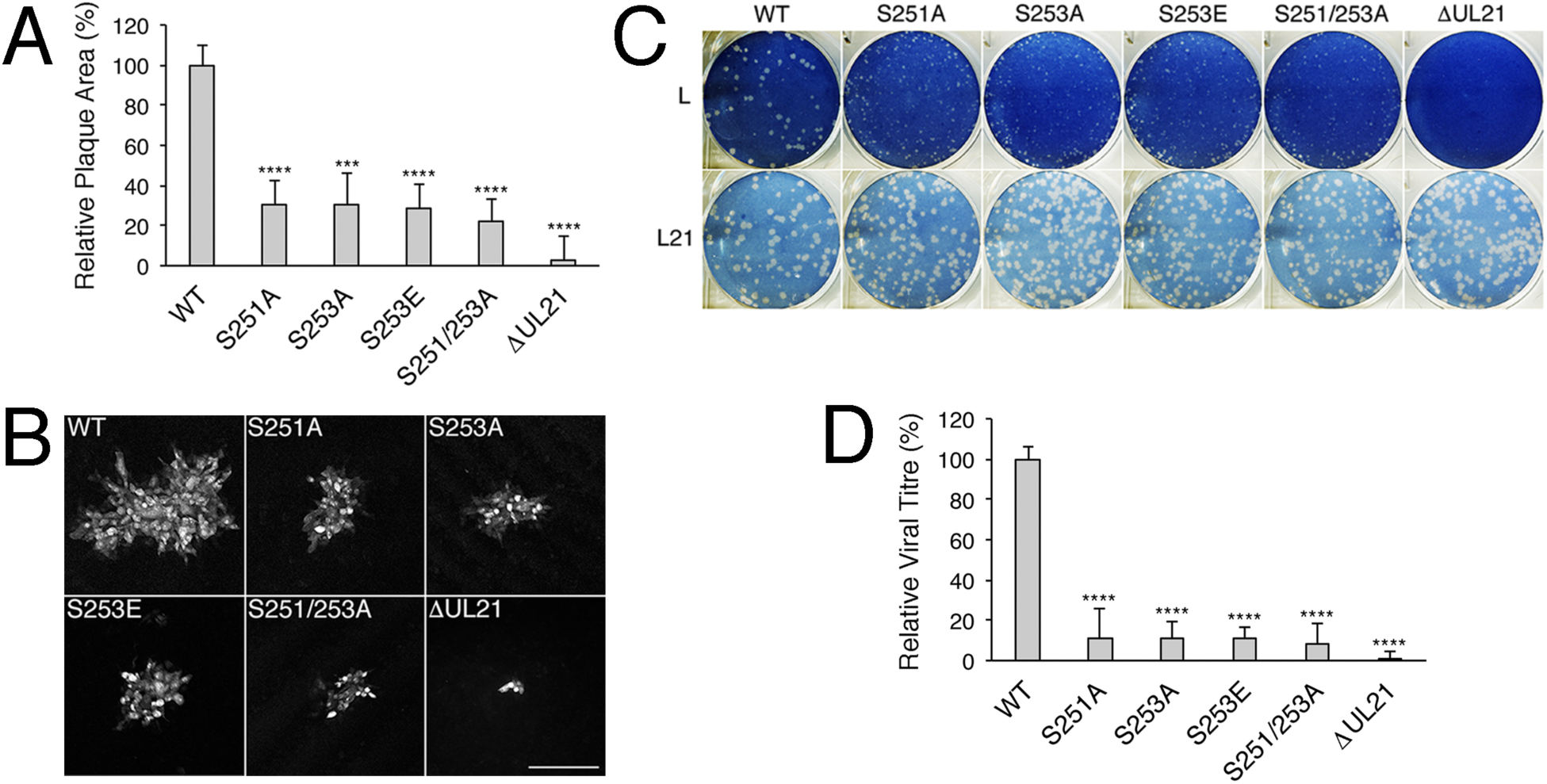
Diminished pUL21 phosphorylation results in reductions in cell-cell spread of virus infection and virus replication. **A.** Vero cell monolayers were infected with the indicated viruses, overlayed with methocel then fixed and stained for pUs3 at 24 hours post infection. The area of 40 plaques per strain were measured using Image Pro software. Plaque areas were scored relative to the area of wild type (WT) plaques. Error bars are standard error of the mean. Asterisks indicate P values: *** ≤ 0.001; **** ≤ 0.0001. **B.** Representative images of plaques formed by the indicated viruses identified by staining with pUs3 antisera. Scale bar is 200 μm. **C.** L or L21 cell monolayers were infected with the indicated viruses, overlayed with medium containing 1% methocel then fixed and stained with 0.5% methylene blue in 70% methanol at 3 days post infection to visualize plaque formation. **D.** Vero cells were infected in triplicate with the indicated viruses at a multiplicity of infection of 0.1 for 48 hours then cells and medium were harvested together. Titres of the harvested material were determined by duplicate plaque assay on L21 cells and scored relative to the titres of WT virus. Error bars are standard error of the mean. Asterisks indicate P values: **** ≤ 0.0001.

### Diminished pUL21 phosphorylation impairs secondary envelopment of cytoplasmic nucleocapsids

To begin investigating what underlies the small plaque phenotype of HSV-2 viruses with defects in pUL21 phosphorylation, we analyzed Vero cells infected with these viruses by TEM. This analysis revealed substantive numbers of nucleocapsids in the cytoplasm indicating that deficiencies in pUL21 phosphorylation had no discernible impact on nuclear egress. In keeping with this view, no major differences were observed in the phosphorylation profile of the NEC component pUL31 in cells infected with pUL21 phosphorylation deficient viruses in comparison to cells infected with WT (Figure 3D). However, most of the cytoplasmic nucleocapsids were not enveloped, indicating a defect in secondary envelopment. To quantify this observation, the ratio of non-enveloped to enveloped cytoplasmic nucleocapsids from micrographs of infected cells was scored (Table 4). The ratio of non-enveloped to enveloped cytoplasmic nucleocapsids in WT HSV-2 strain 186 infected cells was approximately 1 and this value increased 7 to 26-fold in viruses deficient in pUL21 phosphorylation supporting the notion that secondary envelopment was defective in the absence of pUL21 phosphorylation. To contextualize this result, we consulted the literature on the role of pUL21 in secondary envelopment and found that this has only been demonstrated experimentally for bovine herpesvirus 1 (4). It has been speculated that HSV pUL21 plays a role in secondary envelopment (32, 33), however this has not been quantified. To rectify this, we analyzed our previous TEM micrographs of cells infected with WT and pUL21 deficient HSV-2 and HSV-1 strains. The ratio of non-enveloped to enveloped cytoplasmic nucleocapsids was increased 3 to 12-fold over WT parental strains in the case of HSV-2 viruses deficient in pUL21 and 4 to 8-fold over WT parental strains in the case of HSV-1 viruses deficient in pUL21 (Table 4). The pUL21 binding partner, pUL16, has been demonstrated experimentally to play a role in secondary envelopment (34, 35) and similar increases in the ratio of non-enveloped to enveloped cytoplasmic nucleocapsids have been reported for HSV-2 and HSV-1 viruses lacking pUL16 (34).

**Table 4.** Analysis of secondary envelopment in HSV-2 and HSV-1.

| Strain (# of micrographs scored) | Ratio of non-enveloped to enveloped cytoplasmic nucleocapsids (fold increase relative to WT) | Average # of cytoplasmic nucleocapsids per micrograph (total # of cytoplasmic nucleocapsids scored) |
| --- | --- | --- |
| WT HSV-2 186 (12) | 0.8 (1) | 18.8 (225) |
| HSV-2 186 S251A (16) | 5.4 (6.8) | 27.9 (447) |
| HSV-2 186 S253A (12) | 6.7 (8.4) | 24.3 (292) |
| HSV-2 186 S253E (12) | 9.5 (11.9) | 27.1 (325) |
| HSV-2 186 S251/253A (13) | 21 (26.3) | 32.1 (417) |
| HSV-2 186 ΔUL21 (10) | 9.3 (11.6) | 10.3 (103) |
| WT HSV-2 HG52 (13) | 2.7 (1) | 18.7 (243) |
| HSV-2 HG52 ΔUL21 (9) | 13.1 (4.9) | 28.1 (253) |
| WT HSV-2 SD90e (11) | 6.6 (1) | 33.7 (371) |
| HSV-2 SD90e ΔUL21 (12) | 19.3 (2.9) | 45.7(548) |
| WT HSV-1 F (14) | 1.2 (1) | 25.2 (353) |
| HSV-1 F ΔUL21 (12) | 4.2 (3.5) | 25.7 (308) |
| WT HSV-1 KOS (16) | 1.9 (1) | 31.3 (500) |
| HSV-1 KOS ΔUL21 (12) | 14.3 (7.5) | 52.3 (628) |

## Discussion

The data described above indicate that HSV-2 pUL21 is phosphorylated on at least two serines that reside within its linker region. Mutation of either serine 251 or serine 253 of pUL21 resulted in diminished pUL21 phosphorylation, which confirmed the mass spectrometry-based prediction of these phosphorylation sites. As pUL21-derived peptides with multiple phosphorylated residues were never detected in our mass spectrometry analyses and multiple phosphoforms of pUL21 were never detected in our phos-tag analyses, we speculate that only one site in the linker region is phosphorylated at any given time. We note that, in addition to phosphorylation on either serine 251 or serine 253, phosphorylation on serine 261 is hypothetically possible although phosphorylation at this site was only predicted once in our mass spectrometry analyses.

Based on our observation that the phosphorylated form of pUL21 was not detected in cells infected with two independently isolated ΔUL13 strains, we propose that pUL21 is a substrate of pUL13. Direct phosphorylation of pUL21 by pUL13 is the most straightforward explanation for this result, however, the involvement of a cellular kinase that requires pUL13 expression to phosphorylate pUL21 cannot be excluded. Many viral and cellular substrates have been described for pUL13 and its orthologs (reviewed in (36, 37)), including pUL13 itself (38–40). The minimal recognition site for HSV-2 pUL13 has been defined using *in vitro* kinase assays as the amino acid pair serine-proline flanked by alanines and glycines (38). Of the serines identified as potential pUL21 phosphorylation sites in our mass spectrometry analyses, serine 251 best fits this simplistic criterion although it is flanked by valines which, like alanines, are aliphatic amino acids.

The HSV-2 pUL21 validated phosphorylation sites serine 251 and serine 253 and the predicted phosphorylation site serine 261 are proline, glycine, and asparagine, respectively in the linker region of HSV-1 pUL21 (Figure 3A). The amino acid differences between HSV-2 and HSV-1 pUL21 may explain why we were unable to detect phosphorylated pUL21 in cells infected with HSV-1. Interestingly, serines 251 and 253 are both conserved in pUL21 from chimpanzee herpesvirus (ChHV), which is more closely related to HSV-2 than HSV-1 is to HSV-2 (41, 42). It will be of interest in future studies to investigate if differences in pUL21 phosphorylation may underlie our observations that the requirements for pUL21 differ between HSV species (13).

As the defects in cell-cell spread of virus infection and virus replication noted with the phosphomimetic mutant S253E were comparable to that of phosphodeficient mutants S251A, S253A and S251/253A, we hypothesize that reversible phosphorylation is required for HSV-2 pUL21 functionality. Reversible phosphorylation may account for our ability to detect both phosphorylated and non-phosphorylated forms of pUL21 in cells infected with WT HSV-2 at late times post infection. Alternatively, replacement of a single serine with glutamic acid may be insufficient to mimic pUL21 phosphorylation or loss of a serine in this area of the linker may disrupt pUL21 functionality in a manner that is not related to phosphorylation. Future mutational analysis of the linker region of HSV-2 and HSV-1 pUL21 should help distinguish amongst these possibilities.

During HSV virion morphogenesis, nucleocapsids are assembled in the nucleus followed by their translocation across the nuclear envelope and into the cytoplasm where the final stages of virion assembly take place. The process of nuclear egress begins with a nucleocapsid acquiring a primary envelope by budding into the inner nuclear membrane resulting in the formation of a transient perinuclear enveloped virion. The envelope of the perinuclear enveloped virion then fuses with the outer nuclear membrane resulting in the deposition of the nucleocapsid into the cytoplasm. Cytoplasmic nucleocapsids then traffic to membranes derived from the trans-Golgi network, or an endosomal compartment, where secondary envelopment of the nucleocapsid take place resulting in the formation of a mature virion. The importance of pUL21 in nuclear egress is well established (1, 5, 6, 8, 43) and is likely a consequence of pUL21-mediated targeting of PP1α towards the nuclear egress complex components, pUL31 and pUL34, leading to their dephosphorylation and activation (6, 8). While the PP1α binding site in pUL21, like serines 251 and 253, is located within the linker region it is unlikely that PP1α binding to pUL21 is regulated by its phosphorylation as this would be expected to interfere with nuclear egress. The results of our ultrastructure analysis of cells infected with phosphorylation-deficient viruses (Figure 5) suggest that the role of pUL21 in nuclear egress is not dependent on its phosphorylation status.

**Figure 5.**
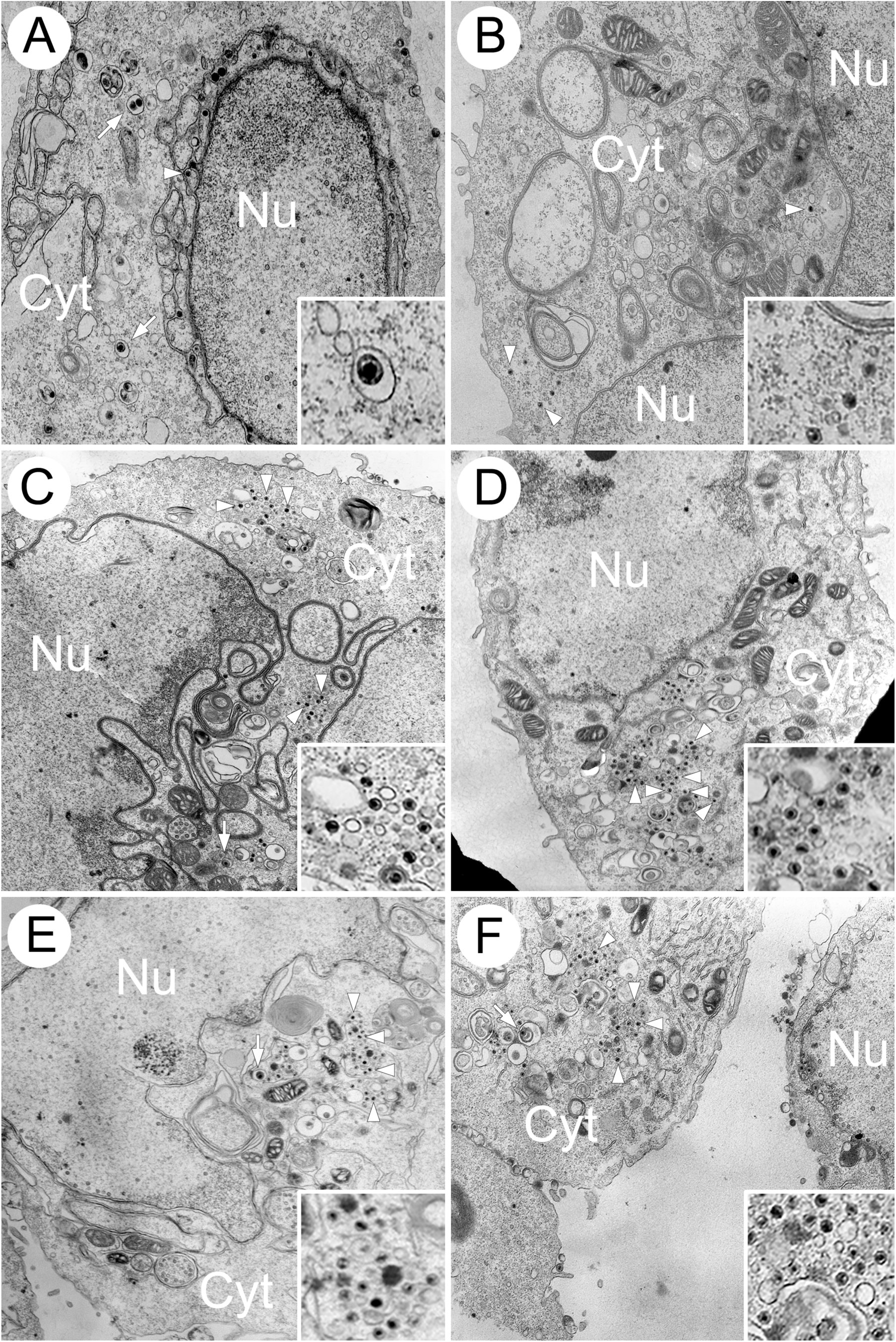
Ultrastructural analysis of cells infected with viruses carrying mutations in pUL21 serine 251 and/or serine 253. Vero cells were infected at a multiplicity of infection of 3. At 18 hours post infection, cells were fixed and processed for TEM as described in Materials and Methods. Representative images of WT (A), 1′UL21 (B), S253E (C), S251A (D), S253A (E), and S251/253A (F) infected cells are shown. Arrows indicate enveloped nucleocapsids and arrowheads indicate non-enveloped nucleocapsids. Nu=nucleus; Cyt=cytoplasm.

Additionally, unlike what was observed in pUL21 deletion strains (6), the phosphorylation profiles of pUL31 in lysates from cells infected with pUL21 phosphorylation-deficient viruses were indistinguishable from WT infected cells (Figure 3D) suggesting that pUL21-mediated regulation of nuclear egress complex components is unperturbed by pUL21 phosphorylation. The importance of pUL21 in secondary envelopment has been demonstrated for bovine herpesvirus (4), however, the importance of HSV pUL21 in secondary envelopment was speculative prior to this study. Our ultrastructure analysis of cells infected with multiple HSV strains lacking pUL21 now confirms the importance of HSV pUL21 in secondary envelopment as well as a requirement for phosphorylated pUL21 in the case of HSV-2. Furthermore, we now have a tool (S251/S253A) for discerning the role of pUL21 in nuclear egress from its role in secondary envelopment.

Finally, as detailed in the Introduction, pUL21 is a multifunctional protein. It is possible that other HSV-2 pUL21 functions could be affected by its phosphorylation status and so additional activities that rely on its phosphorylation status may be revealed. Thus, phosphorylation status should be taken into consideration when studying other roles for pUL21 during infection and/or its ability to interact with viral and cellular proteins.

## Acknowledgements

This work was supported by the Canadian Institutes of Health Research Project Grants 407982 and 486466, Natural Sciences and Engineering Research Council of Canada Discovery Grant RGPIN-2018-04249 and Canadian Foundation for Innovation award 16389 to BWB. JHM was supported in part by a Franklin Bracken Fellowship awarded by Queen’s University. The funders had no role in study design, data collection and interpretation or the decision to submit the work for publication. We thank Dr. Y. Kawaguchi (University of Tokyo), Dr. D. Knipe (Harvard University), Dr. L.W. Enquist (Princeton University), Dr. G.A. Smith (Northwestern University) and Dr. N Osterrieder (Freie Universität Berlin) for providing reagents used in this study. We thank Dr. X. Yan (Queen’s University) for assistance with preparing samples for TEM analyses and Dr. L. Brechenmacher (Southern Alberta Mass Spectrometry Facility, University of Calgary) for expert guidance with mass spectrometry experiments. We are grateful to Dr. S.C. Graham (Cambridge University) and T. Vey (Queen’s University) for valuable input and to members of the Banfield laboratory for helpful comments on the manuscript.

